# scFeatures: Multi-view representations of single-cell and spatial data for disease outcome prediction

**DOI:** 10.1101/2022.01.20.476845

**Authors:** Yue Cao, Yingxin Lin, Ellis Patrick, Pengyi Yang, Jean Yee Hwa Yang

## Abstract

Recent advances in single-cell technologies enable scientists to measure molecular data at high-resolutions and hold the promise to substantially improve clinical outcomes through personalised medicine. However, due to a lack of tools specifically designed to represent each sample (e.g. patient) from the collection of cells sequenced, disease outcome prediction on the sample level remains a challenging task. Here, we present scFeatures, a tool that creates interpretable molecular representation of single-cell and spatial data using 17 types of features motivated by current literature. The feature types span across six distinct categories including cell type proportions, cell type specific gene expressions, cell type specific pathway scores, cell type specific cell–cell interaction scores, overall aggregated gene expressions and spatial metrics. By generating molecular representation using scFeatures for single-cell RNA-seq, spatial proteomic and spatial transcriptomic data, we demonstrate that different types of features are important for predicting different disease outcomes in different datasets and the downstream analysis of features uncover novel biological discoveries.

## Introduction

Recent single-cell or near single-cell resolution omics technologies such as spatial transcriptomics enable the discovery of cell- and cell type-specific knowledge and have transformed our understanding of biological systems, including diseases ^1^. Key to the exploration of such data is the ability to untangle and extract useful information from their high feature dimensions ^2^ and uncover hidden insights. A plethora of computational methods has been developed on this front, with the main focus on individual cell analysis ^3^, such as cell type identity ^4 5^ and pseudotime ordering within a lineage ^6^. While these tools enable characterisation of individual cells, there is a lack of tools that allow for the representation of individual samples based on their cellular characteristics and the investigation of how these cellular properties are driving disease outcomes. With the recent surge of multi-condition and multi-sample single-cell studies on large sample cohort ^7^, the next frontier of research is on representing and characterising cellular properties at the sample (e.g. individual patient) level for linking such information with the disease outcome.

Creating a representation of each sample from the collection of sequenced cells is a crucial step for subsequent analysis as successful modelling and interpretation of disease outcome requires biologically relevant learning features from the data. While using the original expression matrix as the input to various models could inform the change in transcriptomics level across disease conditions, the ability to represent the data with other layers of information is critical for uncovering additional insights given the complex and nonlinear relationships among the feature dimensions (e.g. interaction of genes, gene networks and pathways). The single-cell field has a wealth of tools for data exploration ^8^ which enables exploration of biology underlying the individuals. Most current tools are not specifically designed to derive a set of features that can be used to represent an individual. Yet, with careful adaptation, a number of approaches can be used to construct novel molecular representations of individual samples. Cell-cell interactions tools ^9^, for example, calculate cell type-specific signalling scores between pairs of ligand and receptor molecules. The interaction scores can be used to represent the intercellular communications of cells and cell types in a sample. Another example is gene set enrichment analysis ^10^ which infers the pathway enrichment score of individual cells. By summarising the scores across cell types, a cell type-specific representation of the pathway enrichment of each sample can be constructed.

To this end, we develop scFeatures, a tool that generates a large collection of interpretable molecular representations for individual samples in single-cell omics data, which can be readily used by any machine learning algorithms to perform disease outcome prediction and drive biological discovery. Together, scFeatures generates features across six categories representing different molecular views of cellular characteristics. These include i) cell type proportions, ii) cell type specific gene expressions, iii) cell type specific pathway expressions, iv) cell type specific cell-cell interaction (CCI) scores, v) overall aggregated gene expressions and vi) spatial metrics. The different types of features constructed thereby enables a multi-view of our data and enables a more comprehensive representation of the expression data. In a collection of 17 published single-cell RNA-seq, single-cell spatial proteomics and spatial transcriptomics datasets, scFeatures reveal different feature classes are useful for predicting the disease outcomes in different datasets. Furthermore, through examining the selected features in two case studies, scFeatures uncovers cell types important to ulcerative colitis and stratified patients with distinct survival outcomes in triple negative breast cancer patients. Together, these results demonstrate that scFeature enables a data-driven generation (or feature engineering)and facilitate unbiased identification of feature classes most perturbed by the disease conditions.

## Results

### scFeatures performs multi-view feature engineering for single-cell and spot-based data

We propose scFeatures, a new multi-view feature engineering framework that creates an interpretable representation of cellular level features for each individual sample from a given single-cell or spot-based expression dataset (Fig. 1a). To capture the wide range of cellular information for sample classification (e.g., patients versus healthy individuals) using single-cell data, we implemented an extensive collection of algorithms to extract over 50,000 interpretable features from a given dataset. These features, spanning a total of 17 types, are motivated by established analytical approaches in a broad range single-cell literature and can be broadly grouped into six distinct categories including i) cell type proportions, ii) cell type specific gene expressions, iii) cell type specific pathway expressions, iv) cell type specific CCI scores, v) overall aggregated gene expressions and vi) spatial metrics (Fig. 1b). These collections of constructed features can then be used for various downstream analysis such as disease outcome prediction, biomarker selection, survival analysis and enable the identification of interpretable features and feature types associated with disease conditions.

**Figure 1.**
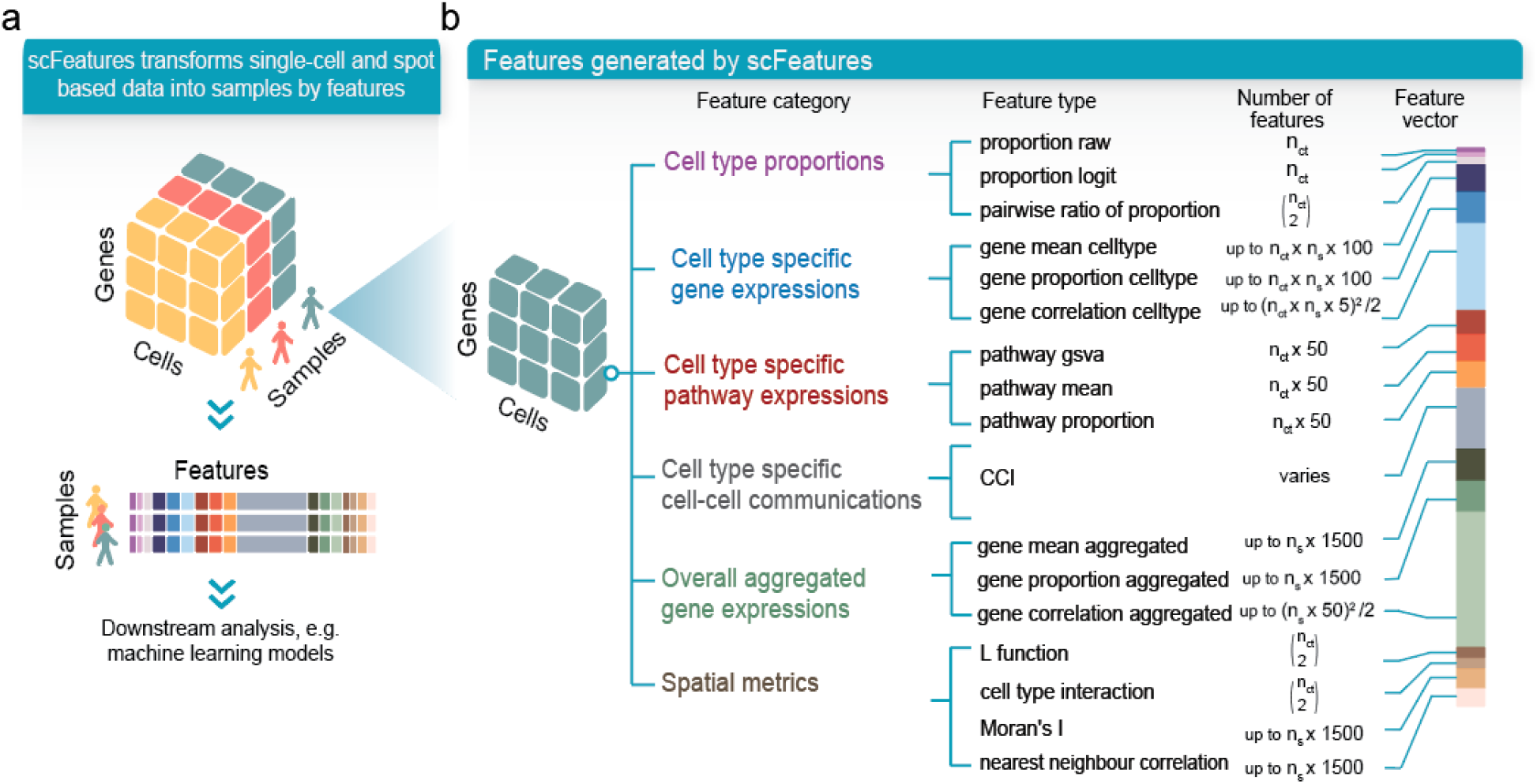
Overview of scFeatures. **a** The input for scFeatures is an omics dataset containing multiple samples such as patients. scFeatures extracts different views of the data, thereby transforming the gene by cell matrix into a vector of features for each sample. **b** scFeatures constructs 17 feature types that can be broadly classified into six categories. Each feature type consists of multiple individual features. For example, for “gene mean cell type”, 100 features are generated by default per cell type (n_ct_) per sample (n_s_) (see Methods).

The six feature categories represent different “views” of the single-cell information. Specifically, category I captures cell type proportion information in which the proportion of cell types for each sample and the ratio of proportions between two cell types are measured. Category II represents cell type specific gene expression, and examines the expression of sets of genes or proteins in each cell type. We implemented different approaches for representing genes or proteins measurement, including average expression, proportion of expression and correlation of expressions. In category III, which calculates cell type specific pathway scores, by default the 50 hallmark pathways in the Molecular Signatures Database (MSigDB) ^11 12^ were used to generate various features such as the average expression of each pathway in each cell type. Category IV contains the CCI scores, measuring the probability of ligand-receptor interaction based on the expression values of each sample. Category V is designed to recreate the bulk expression through aggregating the expression across cells or spots depending on the data types. Category VI is designed specifically for spatial data type for capturing spatial information and includes classical metrics for identifying spatial patterns.

scFeatures extracts interpretable features from data generated by scRNA-seq, spatial proteomics, and spatial transcriptomics (Table 1). In particular, spatial transcriptomics data, is a spot-based technique in which the expression value of each spot is based on a small population of cells often containing cells from multiple cell types. We developed several novel ways to adapt the 13 feature types to spot-based data whenever possible; this collection of spatial metrics takes into consideration the properties of spot-based technology and reveals cell typespecific features in spot-based data. For example, spot-based data precludes direct application of cell type proportion computation since each spot includes an unknown number of cells while cell type percentage estimation requires individual cell counts for each cell type. To overcome this issue, we estimated the number of cells in each spot using the library size of that location, based on the association between the two values. Supplementary Table 1 provides more documentation on the implementation details on the adaptation of feature types from single cell RNA-sequencing to spot-based technologies.

**Table 1.** List of features generated by scFeatures.

| Applicati<br>on | Data<br>character<br>istic and<br>represent<br>ative<br>technolo<br>gy | Featur<br>e<br>catego<br>ry | Cell type specific<br>proportions |  |  | Cell type specific gene<br>expressions |  |  | Cell type specific<br>pathway<br>expressions |  |  | Cell -<br>cell<br>interac<br>tion<br>scores | Overall aggregated gene<br>expressions |  |  | Spatial metrics |  |  |  |
| --- | --- | --- | --- | --- | --- | --- | --- | --- | --- | --- | --- | --- | --- | --- | --- | --- | --- | --- | --- |
|  |  |  | Pro<br>porti<br>on<br>raw | Proport<br>ion<br>logit | Rati<br>o | Mea<br>n | Proport<br>ion | Correla<br>tion | Mea<br>n | GS<br>VA | Pro<br>porti<br>on |  | Mea<br>n | Proport<br>ion | Correla<br>tion | L<br>func<br>tion | Cell<br>type<br>inter<br>acti<br>on | Mor<br>an's<br>I | Nearest<br>neighbour<br>correlatio<br>n |
| scRNA-seq data | Single-cell<br>based<br>Eg. 10x,<br>Smart-Seq |  | ✓ | ✓ | ✓ | ✓ | ✓ | ✓ | ✓ | ✓ | ✓ | ✓ | ✓ | ✓ | ✓ |  |  |  |  |
| Spatial<br>proteomic<br>s data | Single-cell<br>based<br>Eg. MIBI-TOF,<br>IMC,<br>CODEX |  | ✓ | ✓ | ✓ | ✓ | ✓ | ✓ |  |  |  |  | ✓ | ✓ | ✓ | ✓ | ✓ | ✓ | ✓ |
| Single cell<br>transcript<br>ome data<br>data | Spot<br>based<br>Eg. ST,<br>Visium,<br>MALDI |  | ✓ | ✓ | ✓ | ✓ |  |  | ✓ |  |  |  | ✓ | ✓ | ✓ | ✓ | ✓ | ✓ | ✓ |

### scFeatures generates large collection of diverse features and is scalable to large datasets

To demonstrate the characteristics of the feature representation, we applied scFeatures to 17 datasets measured using scRNA-seq, spatial proteomics and spatial transcriptomics data (Table 2). For a typical scRNA-seq data, scFeatures generated over 50,000 features (Fig. 2a). As expected, the number of features generated were mostly associated with the number of cell types in the dataset and not other data characteristics such as number of genes, number of cells and number of patients (Supplementary Figure 1).

**Figure 2.**
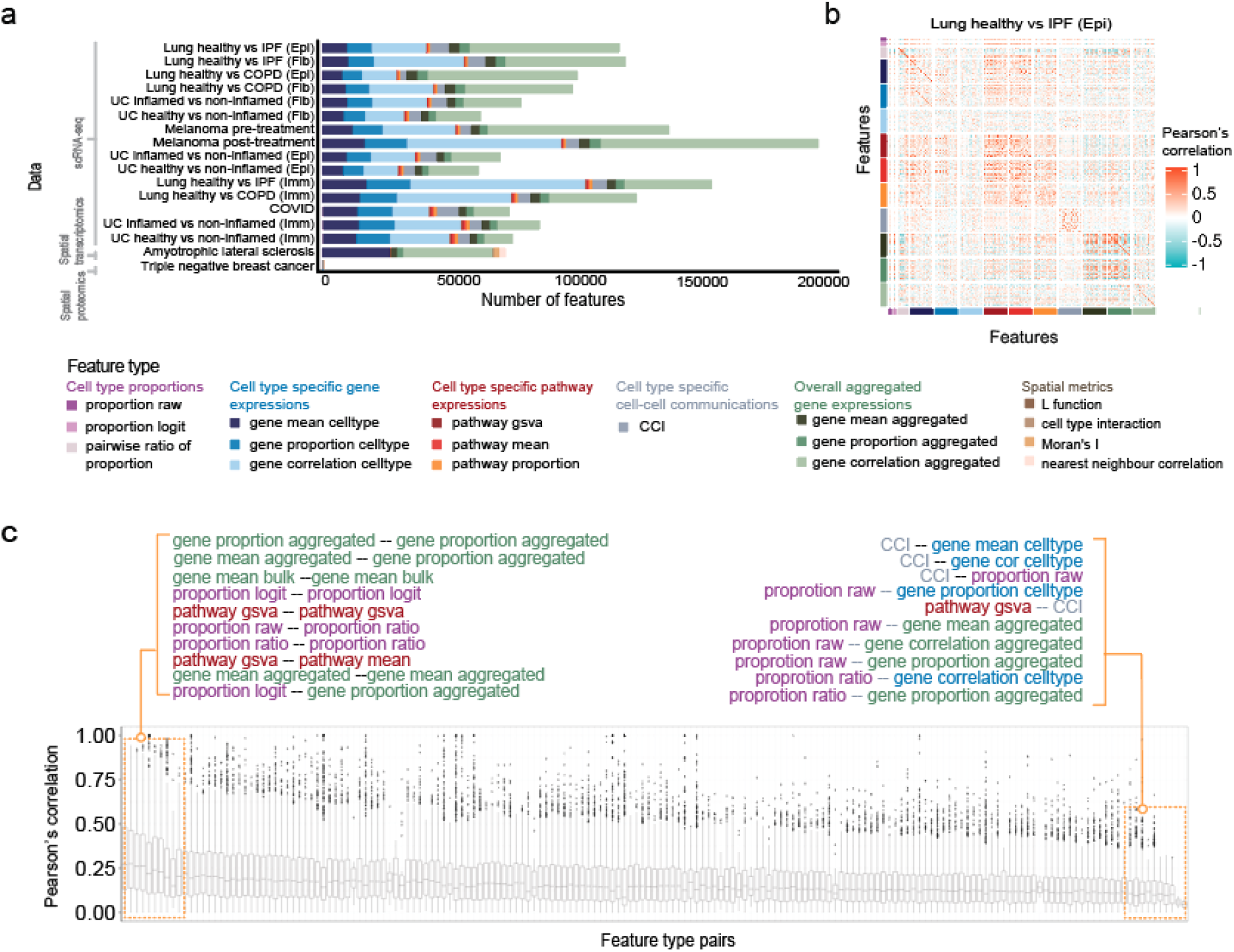
Characteristics of the features generated by scFeatures. **a** Compositional barchart showing the number of features generated by scFeatures for each dataset. Datasets are first ordered by data types, and then by the number of cell types. **b** Correlation plot showing Pearson’s correlation of features on the “Lung healthy vs IPF (Epi)” dataset as a representative example. The features are colour labelled by feature types for ease of interpretation. **c** Boxplots summarising the correlation between pairs of features across all datasets (see Methods). Texts highlight the 10 most and 10 least correlated feature types pairs.

**Table 2.** Details of the datasets used in the study.

| Dataset name referred in the study | Study reference | Outcome | Number of genes/proteins | Number of cells/spots | Number of samples | Species | Type of data |
| --- | --- | --- | --- | --- | --- | --- | --- |
| Lung healthy vs IPF (Epi) | Adams et al. 2020<sup>20</sup> | Healthy vs IPF | 45947 | 17970 | 53 | Human | scRNA-seq |
| Lung healthy vs IPF (Fib) |  | Healthy vs IPF | 45947 | 12753 | 49 | Human | scRNA-seq |
| Lung healthy vs IPF (Imm) |  | Healthy vs IPF | 45947 | 208774 | 60 | Human | scRNA-seq |
| Lung healthy vs COPD (Epi) |  | Healthy vs COPD | 45947 | 7888 | 38 | Human | scRNA-seq |
| Lung healthy vs COPD (Fib) |  | Healthy vs COPD | 45947 | 5286 | 36 | Human | scRNA-seq |
| Lung healthy vs COPD (Imm) |  | Healthy vs COPD | 45947 | 149875 | 46 | Human | scRNA-seq |
| UC healthy vs non-inflamed (Epi) | Smillie et al. 2019<sup>13</sup> | Healthy vs non-inflamed | 20028 | 99962 | 30 | Human | scRNA-seq |
| UC healthy vs non-inflamed (Fib) |  | Healthy vs non-inflamed | 19076 | 21627 | 30 | Human | scRNA-seq |
| UC healthy vs non-inflamed (Imm) |  | Healthy vs non-inflamed | 19076 | 118784 | 30 | Human | scRNA-seq |
| UC inflamed vs non-inflamed (Epi) |  | Inflamed vs non-inflamed | 20028 | 72748 | 35 | Human | scRNA-seq |
| UC inflamed vs non-inflamed (Fib) |  | Inflamed vs non-inflamed | 19076 | 23392 | 36 | Human | scRNA-seq |
| UC inflamed vs non-inflamed (Imm) |  | Inflamed vs non-inflamed | 20529 | 159242 | 36 | Human | scRNA-seq |
| Melanoma pre-treatment | Sade-Feldman et al. 2018 <sup>21</sup> | Responder vs non-responder | 50513 | 5925 | 19 | Human | scRNA-seq |
| Melanoma post-treatment |  | Responder vs non-responder | 50513 | 10357 | 29 | Human | scRNA-seq |
| COVID | Schulte-Schrepping et al. 2020 <sup>22</sup> | Mild vs severe | 24794 | 48069 | 27 | Human | scRNA-seq |
| Amyotrophic lateral sclerosis | Maniatis et al. 2019 <sup>24</sup> | ALS vs normal | 9129 | 23373 | 33 | Mouse | Spatial transcriptomics |
| Triple negative breast cancer | Keren et al. 2019 <sup>28</sup> | Survival period | 38 | 199817 | 39 | Human | Spatial proteomics |

To explore the complementarity among the features generated from scFeatures, we examined the correlation between the features across 17 datasets. We observed a mosaic pattern, where feature types from the same feature category tended to have higher correlations with each other but very low correlation with feature types from other categories (Fig. 2b and Supplementary Figure 2). By summarising the correlation values between every pairwise combination of feature types (Supplementary Figure 3), we found that, overall, the feature types were poorly correlated (Fig. 2c) with the median correlation between feature types ranges between 0.1 to 0.25 (Fig. 2c). The relatively low correlations indicate that the features constructed by scFeatures were diverse and potentially complementary for exploring disease outcomes. The top three correlated pairs were observed amongst feature types related to overall aggregated gene expression. This is consistent with our expectation of some degree of co-expression linked with various conditions (e.g., healthy or disease outcome), and it is important to emphasise that these top correlated features remain poorly correlated.

We next benchmarked the runtime and memory requirement of the feature types on single-cell scRNA-seq (Supplementary Figure 4a), spatial proteomics (Supplementary Figure 4b), as well as on spot-based spatial transcriptomics datasets (Supplementary Figure 4c) for evaluating both the single-cell RNA-sequencing implementation and the spot-based implementation. All datasets contain 1,000 to 100,000 cells. On the largest datasets with 100,000 cells, the majority of feature types took less than a minute to compute when executed on eight cores, demonstrating that scFeatures is highly scalable to large datasets. As expected, there was some trade-off between processing time and memory. As a result of parallel computation over eight cores, some feature types required more than 10GB of RAM in total; however, users can run on a single core to decrease the memory required.

### The most informative features classes differ between different datasets

We hypothesised that distinct feature classes would be informative for different datasets since each dataset comprises samples with varying characteristics and disease outcomes. Several datasets were used where each feature type was evaluated on their ability to predict disease outcomes and the observations are in alignment with our hypothesis. First, we used a lung disease dataset collection where the cells were split into the epithelial, immune and fibroblast subset and the outcome of interest was to classify the patients into healthy or idiopathic pulmonary fibrosis (IPF). In Fig. 3a, we visualised the performance classification of the feature types on the three subsets and ordered the feature types according to their performance in the epithelial subsets. This reveals that feature types that achieved the highest accuracy in the epithelial subset are related to cell type proportions (i.e., “proportion ratio”, “proportion logit” and “proportion raw”) (Fig. 3a). In contrast, the performance of feature types on the immune and fibroblast subset clearly does not follow the same trend as on the epithelial subset, demonstrating that different feature types are useful to the three datasets

**Figure 3.**
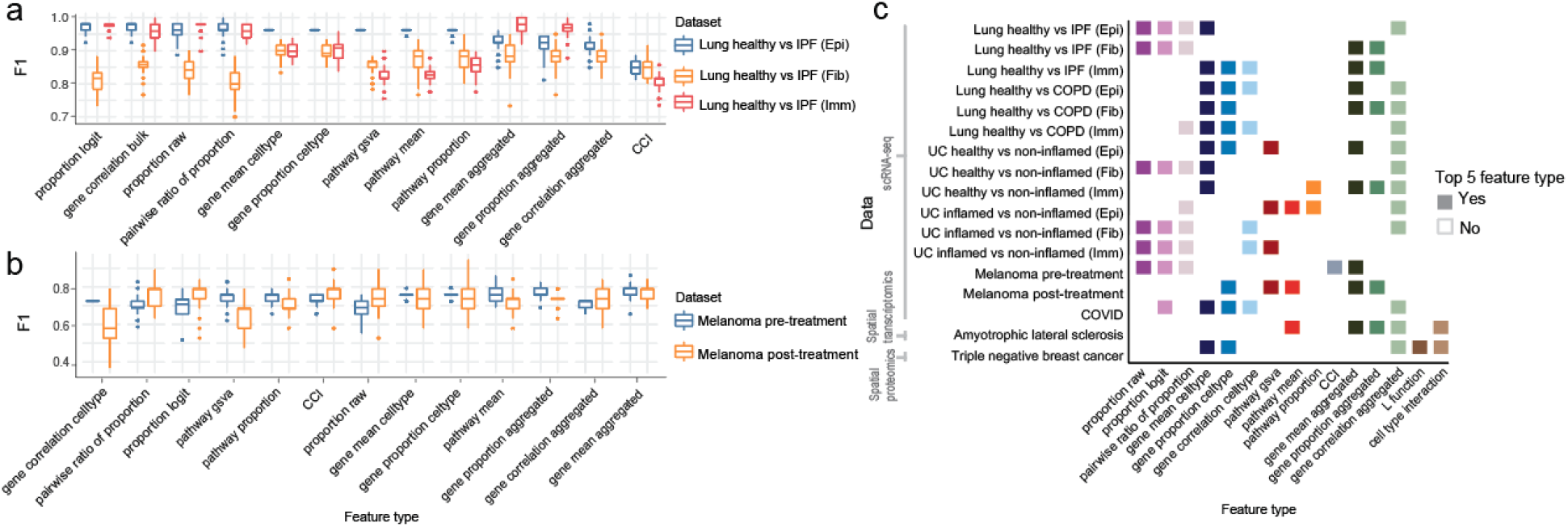
Performance of feature types on patient outcomes. **a** The boxplots compare the F1 score of the feature types on two data collections. Top shows the epithelial, fibroblast and immune subsets of healthy and IPF patients, where the outcome of interest is classifying healthy and IPF status. The feature types are ordered by their F1 scores on the epithelial subst. Bottom shows pre-treatment and post-treatment melanoma patients, where the outcome of interest is classifying therapy responders and non-responders. The feature types are ordered by the difference of the F1 scores between the two datasets. **b** For each dataset, the squares denote the top five feature types with the highest F1 scores.

Similar observation is also found in the melanoma pre-treatment dataset and melanoma posttreatment dataset where the question of interest is classifying non-responder and responders. Fig. 3b illustrate that proportion features (i.e., “proportion raw” and “proportion logit”) more accurately classified patients in the post-treatment dataset than patients in the pre-treatment dataset and pathway features (i.e, “pathway gsva” and “pathway proportion”) provides more accurately classified pre-treated patients.

We then examined across 17 datasets (Supplementary Figure 5) and highlighted the five informative feature types for each dataset (Fig. 3c) for a more comprehensive assessment of the performance of the feature types. Across the 17 datasets tested, “gene mean celltype”, which examines expression in cell type specific manner, occurred in 10 datasets as the top five informative feature types. This is perhaps not surprising, as it elucidates the power of single-cell technology and the ability of cell type specific gene expression to uncover changes in response to diseases. Across the spatial datasets, we saw feature types in the spatial feature category appearing as the top five informative feature types, indicating the usefulness of this category for capturing spatial information and the potential of spatial data modality offering complementary information. All together, these findings highlight that different feature types are useful for exploring disease mechanisms in different datasets and even in different subsets of the same dataset, as seen by the pre- and post-treatment melanoma patients and the lung disease dataset subset by cell types, and argue for the need for a diverse compendium of feature types for such analyses.

### scFeatures provides interpretable insight into disease outcome from scRNA-seq data

To illustrate that scFeatures provides interpretable features for disease understanding, we applied scFeatures on the “UC healthy vs non - inflamed (Fib)” dataset ^13^. This scRNA-seq dataset compares fibroblast cells of non-inflamed biopsies from ulcerative colitis (UC) patients and biopsies from healthy patients. We focused on the two top performing feature types based on the classification model performance from the previous section of “gene mean celltype” and “cell type proportion raw” (Fig. 3c) and discovered different sets of cell types were important to the two feature types. For the feature type based on cell type specific gene expression (denoted by “gene mean celltype”), the top four cell types according to feature importance score (see Methods) are all sub-cell types from WNT2B+ and WNT5B+ cells (Fig. 4a). This indicates that selected features, i.e., genes, from these cell types were considered more important at predicting disease outcomes than genes of other cell types. In contrast, the WNT2B+ and WNT5B+ cell types were ranked as the bottom four cell types in terms of the differences in cell type proportion (in “cell type proportion raw”), indicating that while the gene expression is different between disease outcome, the proportion of cell types are similar. Instead, pericytes, glia, microvascular, and inflammatory fibroblasts were found to be the top four cell types that exhibit differential proportion between healthy and non-inflamed biopsies (Supplementary Figure 6). These two feature types offer different perspectives from the same data and reveal distinct collections of cell types where one group is more concerned with changes in expression and the other collection is more concerned with changes in proportion. It would have been challenging or impossible to accurately disentangle the contributions of cell type percentage and cell specific gene expression in classical bulk gene expression data. These observations not only highlights the necessity of single-cell research, but it also emphasises the significance of evaluating various feature types, as generated by scFeatures.

**Figure 4.**
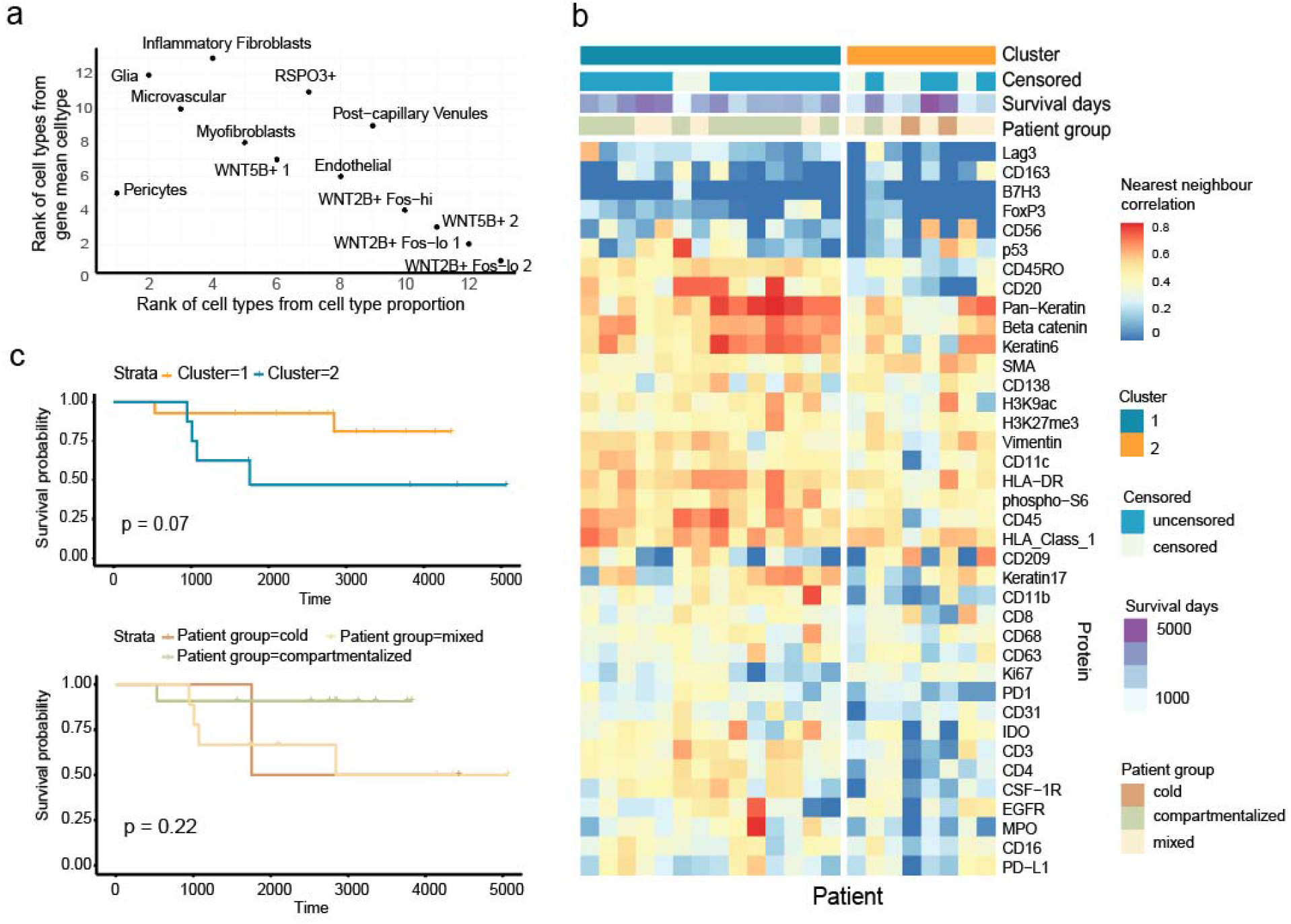
Selected features generated on the “UC healthy vs non - inflamed (Fib)” dataset and the “triple negative breast cancer” datasets. **a** Scatterplot of cell types rank for the feature type “cell type proportion” and “gene mean celltype”. **b** Heatmap showing the clustering result using the nearest neighbour correlation. **c** Kaplan-Meier plot of patients stratified by the clustering output (top) and stratified by patient groups defined in the original study (bottom).

### scFeatures uncovers data features associated with survival outcome from spatial proteomics

To demonstrate the utility of scFeatures at extracting spatial information, we applied scFeatures to a spatial proteomics dataset of tumors from triple negative breast cancer patients. The question of interest is classifying tumors based on cellular organisation into distinct types that are associated with patient survival. The original study defined three tumor groups based on mixing scores, where a “cold group” is identified by low immune infilture, a “compartmentalised group” is identified by compartments formed by almost entirely of either tumour or immune cells, and a “mixed group” is when there is no clear boundary separating the tumor and immune cells.

The nearest neighbour correlation is a feature type in scFeatures that was created primarily to capture spatial co-expression patterns. It computes the correlation of a cell’s protein expression with that of its nearest neighbour. Therefore, spatial organisation of cells, such as whether tumor cells are next to immune cells would affect the correlation of protein expression of cells with neighbouring cells. To construct this feature type, we used scFeatures on selected “triple negative breast cancer” samples from the dataset and clustered the resulting features (Fig. 4b). Survival analysis using the Kaplan-Meier Curve revealed differences between survival outcomes of individuals from the two clusters (P-value of 0.07, Fig. 4c), compared to the patient group defined in the original study with P-value of 0.22. This suggests that the new patient subgroup found by scFeatures has greater association with the survival outcomes and demonstrates the ability of the spatial feature category at representing spatial organisations and uncovering novel patterns in the data.

## Discussion and Conclusion

In summary, scFeatures creates a multi-view molecular representation of individuals by generating over tens of thousands of interpretable features based on single-cell and spot-based spatial data. The innovation and motivation scFeatures lies in the generation of various literature motivated and biologically relevant feature vectors for phenotype disease modelling and disease prediction. We have designed 17 feature types across six categories based on a broad range of analytical approaches in literature from cell type specific gene expression to measures of cellcell (ligand-receptor co-expression) interaction and demonstrated that the feature types are diverse with low correlation amongst them. We demonstrated scFeatures on scRNA-seq data from ulcerative colitis and discovered a number of features linked with disease characteristics and extracting spatial features from a triple negative breast cancer proteomics data resulted in the stratification of tumours that are more strongly related with survival outcomes than the original study’s subgroups.

The features vector generated by scFeatures can be used for a broader set of downstream applications and not limited to the ones illustrated in the case studies. For example, given the features vectors are generated at the sample level, this opens the opportunity for the exploration of differential patient response to diseases due to heterogeneity between individuals. Even for patients recorded as responders to a treatment, the extent of response and the change at omics level varies between individuals. The feature vector can be subjected to latent class analysis, which has typically been applied on single-cell level for exploring cellular diversity ^15, 16^, to enable detection of sub-populations in the cohort, as well as the biology driving patient heterogeneity.

The multiple feature representations generated by scFeatures can be considered as multiple views of the data and as such leads naturally to multi-view learning. This is one of the many collections of methods that perform integration across multiple features classes to enhance model performance. There exists a number of approaches for performing data integration ^17^, from the simple concatenation of features from all feature types into a single vector as the input, to incorporating and optimising the data integration procedure within the model training process. While current multi-view learning in bioinformatics typically refers to the use of multiple omics obtained from the same sample ^18^, we envisage the generation of multiple features types by scFeatures opens new opportunities for multi-view learning from single omic type.

scFeatures is currently designed to perform feature engineering for single-cell RNA-seq, spatial proteomics and spatial transcriptomics data, but the framework is not limited to these platforms. Taking chromatin accessibility as an example, a commonly used analysis strategy is assigning genes based on nearby peaks, thereby converting the peak matrix to a matrix of gene activity scores similar to gene expressions ^19^. Using this approach, all feature classes designed for scRNA-seq are then applicable to chromatin accessibility data. In future, we plan to extend scFeatures to other single-cell omics such as single-cell DNA methylation, single-cell chromatin accessibility and single-cell genomics, leveraging the common analytical approach in these areas and construct specific feature classes. For chromatin accessibility, the co-accessibility between pairs of peaks, which is used to predict cis-regulatory interactions, can be constructed and stored as a vector for each sample. The correlation values between transcription factors (TF) motifs can be readily constructed as another class of feature representation vector, and can be used to identify the modules of TF motifs affected in disease state.

With the recent surge of cohort based single-cell studies and the number of tools for characterising individual cells, there is an increased demand for defining samples in a study based on their cellular characterization to guide better understanding of disease and health. Here, we presented scFeatures, a tool that provides a multi-view extraction of molecular features from single-cell and spot-based spatial data to characterize cellular features of each individual. scFeatures efficiently extracts collections of interpretable features from large-scale data and we demonstrated its ability to derive biological insights in scRNA-seq and spatial data. We envision that scFeatures, a public R package available at https://github.com/SydneyBioX/scFeatures, will facilitate better understanding of single-cell data from a sample (i.e. patient) perspective and the signatures underlying disease conditions from different angles.

## Methods

### Data collection and processing

#### scRNA-seq

To demonstrate scFeatures on scRNA-seq data, we collected data from four published studies and curated a total of 15 datasets from the studies. The data are described in detail below:

##### Six Ulcerative Colitis datasets

The UC data ^13^ sequenced healthy control, inflamed and noninflamed colon biopsies from multiple patients. The data was retrieved from Single Cell Portal with accession ID SCP259. We subset the data into epithelial, stromal cells and immune subset according to the original publication, resulting in the following 6 datasets:

- UC healthy vs non-inflamed (Epi)
- UC healthy vs non-inflamed (Fib)
- UC healthy vs non-inflamed (Imm)
- UC inflamed vs non-inflamed (Epi)
- UC inflamed vs non-inflamed (Fib)
- UC inflamed vs non-inflamed (Imm)

where Epi stands for epithelial, Fib stands for stromal and Imm stands for immune subsets. Inflamed, non-inflamed and healthy are patients’ conditions of interest.

##### Six Lung datasets

The lung data^20^ sequenced healthy control, idiopathic pulmonary fibrosis (IPF), and chronic obstructive pulmonary disease (COPD) biopsies from multiple patients. The data was retrieved from Gene Expression Omnibus (GEO) with accession ID GSE136831. We subset the data into epithelial, stromal cells and immune subset according to the original publication, resulting in the following datasets:

- Lung healthy vs IPF (Epi)
- Lung healthy vs IPF (Fib)
- Lung healthy vs IPF (Imm)
- Lung healthy vs COPD (Epi)
- Lung healthy vs COPD (Fib)
- Lung healthy vs COPD (Imm)

where healthy, IPF and COPD are patients’ conditions of interest.

##### Two melanoma data

^21^ sequenced immune cells from tumor biopsies of melanoma patients prior to and after treatment with immune checkpoint therapy. The data was retrieved from GEO with accession ID GSE120575. We subset the data into pre-treatment and post-treatment datasets. The patients’ conditions of interest in both datasets are non-responding and responding.

##### The COVID dataset

^22^ sequenced peripheral blood mononuclear cells (PBMC) from COVID-19 patients. The data was retrieved from European Genome-phenome Archive (EGA) with accession ID EGAS00001004571. We subset the original data into mild and severe patients and consider the mild and severe disease stage as the patients’ conditions of interest.

### Spatial proteomics

#### The triple negative breast cancer dataset

^23^ measured the patient’s protein expression using MIBI-TOF (multiplexed ion beam imaging by time of flight) technology. Data was obtained from https://mibi-share.ionpath.com.

### Spatial transcriptomics

#### The amyotrophic lateral sclerosis dataset

^24^ sequenced lumbar spinal cord tissue of ALS and control mouse at varying time points using the spatial transcriptomics technology. The data was retrieved from GEO with accession ID GSE120374. We used the subset of data sequenced at the disease onset time point.

### Implementation of feature types

We generated 17 feature types that can be broadly categorised into six categories: i) cell type proportions, ii) cell type specific gene expressions, iii) cell type specific pathway expressions, iv) cell type specific CCI scores, v) overall aggregated gene expressions and vi) spatial metrics. All feature types except for the overall aggregated gene expressions category have different implementations for scRNA-seq and spatial data to better leverage the characteristics of different data types and the implementation details are described in Supplementary Table 1.

For spot-based spatial transcriptomics, we performed the following additional processing in order to allow certain feature classes to be applicable. First, since the cell type specific feature categories require cell type information while the spot in spot-based data contains a mixed population of multiple cells, we used Seurat’s TransferData function to predict the cell type probability of each spot. A published scRNA-seq data on mouse spinal cord with cell type labels was used as the reference ^25^. Then, given that each spot contains an unknown number of cells which vary across each spot, we weighted the contribution of each spot to the generated features by the relative number of cells it contains. We used library size as an estimate of the relative number of cells, motivated by a study that found a high correlation between the number of cells and library size of spots ^26^. To calculate the relative number of cells, we binned the log2 transformed total library size of cells into 100 bins, and assigned each spot a relative number of cells ranging between 1 to 100 according to its bin.

### Correlation between features and feature classes

Given scFeatures constructs a standard matrix of samples by features, we can readily compute the Pearson’s correlation between individual features. In detail, we first computed the correlation between individual features. The median correlation between pairs of feature classes was calculated by taking the absolute values of the individual correlation values of the features in the given pair of feature classes and then computing the median. For the correlation plot shown in Fig. 2b, we subsampled 100 features from feature classes that have more than 100 features to avoid the correlation plot being dominated by feature classes with more features.

### Classification and survival analysis using generated features

In scFeatures, we provide functionality to perform classification and survival analysis for the convenience of users. The classification function is a wrapper around a classification package classifyR^27^ that was published by our group earlier. By default, we use a random forest model, set the number of folds to three, perform 20 cross-validation and calculate F1-score. These were also the settings used to report the classification performance in this study and can be specified by the user. The only exception being that 100 cross-validation was performed to obtain a more stable feature importance score for the case study on the “UC healthy vs noninflamed (Fib)” dataset.

For survival analysis, we use a cox proportional-hazards model provided in the rms R package. By default, we set the number of folds to three, perform 20 cross-validation and calculate C-index. Note that as the cox model is not designed to take in a large number of features at once, unlike a typical classification model, we input one feature from the generated feature class at a time for building the cox model. The best C-index is reported as the performance for the feature class.

### Feature importance score

The runTests function in ClassifyR outputs the features selected by the classification model. Since repeated cross-validation was performed, this generated one set of included features for each cross-validation process. Based on all the derived sets, the frequency of inclusion was considered as the feature importance score of each feature.

For the cell type specific feature category, given that each feature is associated with a cell type, it is also of interest to aggregate the feature importance score associated with each cell type. We approached by summing the feature importance score of all features associated with a cell type, then dividing by the number of features constructed for that particular cell type to adjust for the difference in the number of features per cell type. The final score was considered as the feature importance score of each cell type.

### Speed and memory usage

To benchmark the scalability of the 17 features classes, we used the UC inflamed vs noninflamed (Imm) dataset and took random samples to construct datasets with 1000, 2000, 3000, 5000, 10000, 20000, 30000, 50000, 70000 and 100000 cells. Each dataset contains the same 15 patients and the same 15 cell types.

For the purpose of evaluating the features classes designed for spot-based data which require each spot to be associated with a cell type probability vector, we treated each cell as a “spot” and randomly created a cell type probability vector for each cell. Similarly, for the purpose of evaluating the feature classes under the category spatial metrics which require spatial coordinates of each cell, we randomly assigned a pair of x and y-coordinates to each cell. In addition, the cell type probability and number of cells in each spot was randomly generated to represent such data.

Runtime was measured using the built-in Sys.time function in R. Memory was measured by recording the peak resident set size, which measures the peak amount of memory that a process consumes across all cores. All code was run in parallel using 8 cores for three times and the average measurements were taken. All processes were carried out using a research server with dual Intel(R) Xeon(R) Gold 6148 Processor with 40 cores and 768 GB of memory.

## Supplementary Materials

### Supplementary Figures

**Supplementary Figure 1.**
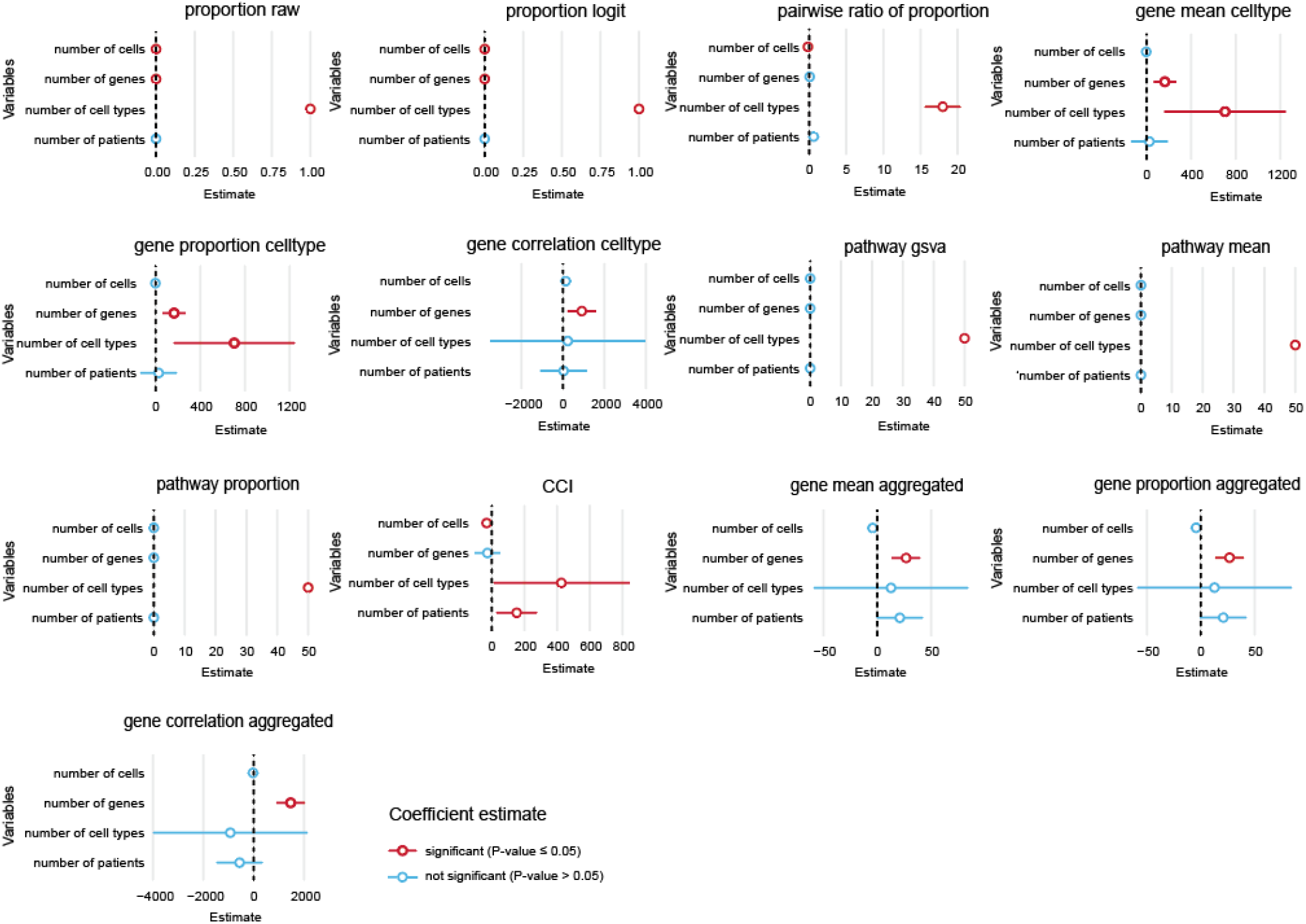
Impact of dataset characteristics on number of features generated by scFeatures. We generated features on 15 scRNA-seq datasets (see Methods). Linear regression model wa**s** fitted to explore the relationship between the number of features and dataset characteristics such as number of cells, genes, cell types and patients. The regression coefficient for each variable is shown in the line plots, with red denoting a significant relationship and blue denoting an insignificant relationship.

**Supplementary Figure 2.**
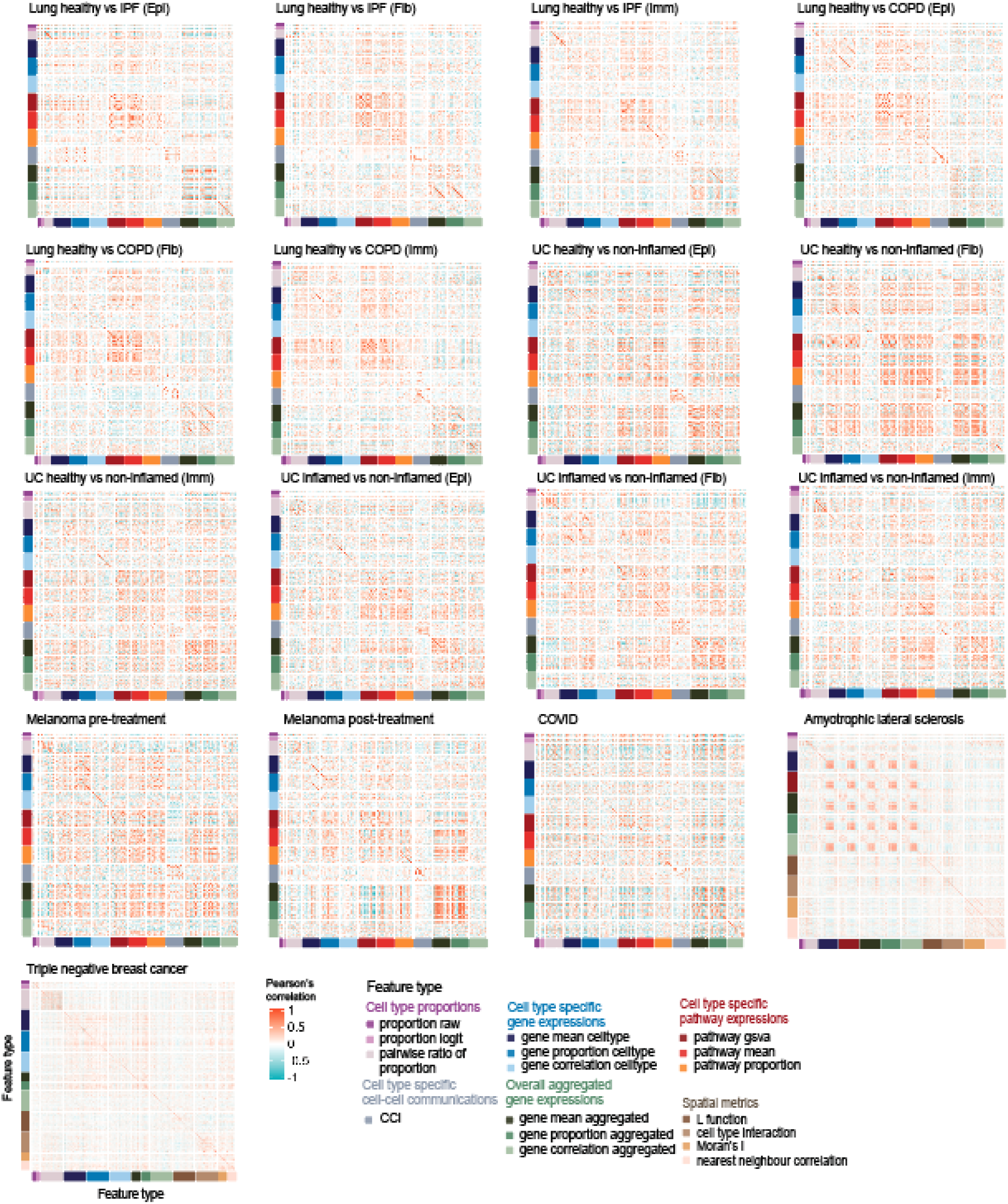
Correlation amongst feature pairs for each dataset. Plots show the Pearson’s correlation between features on each dataset. The features are colour labelled by feature class for ease of interpretation. To avoid the correlation plot being dominated by feature classes with more features, we subsampled 100 features from feature classes with more than 100 features.

**Supplementary Figure 3.**
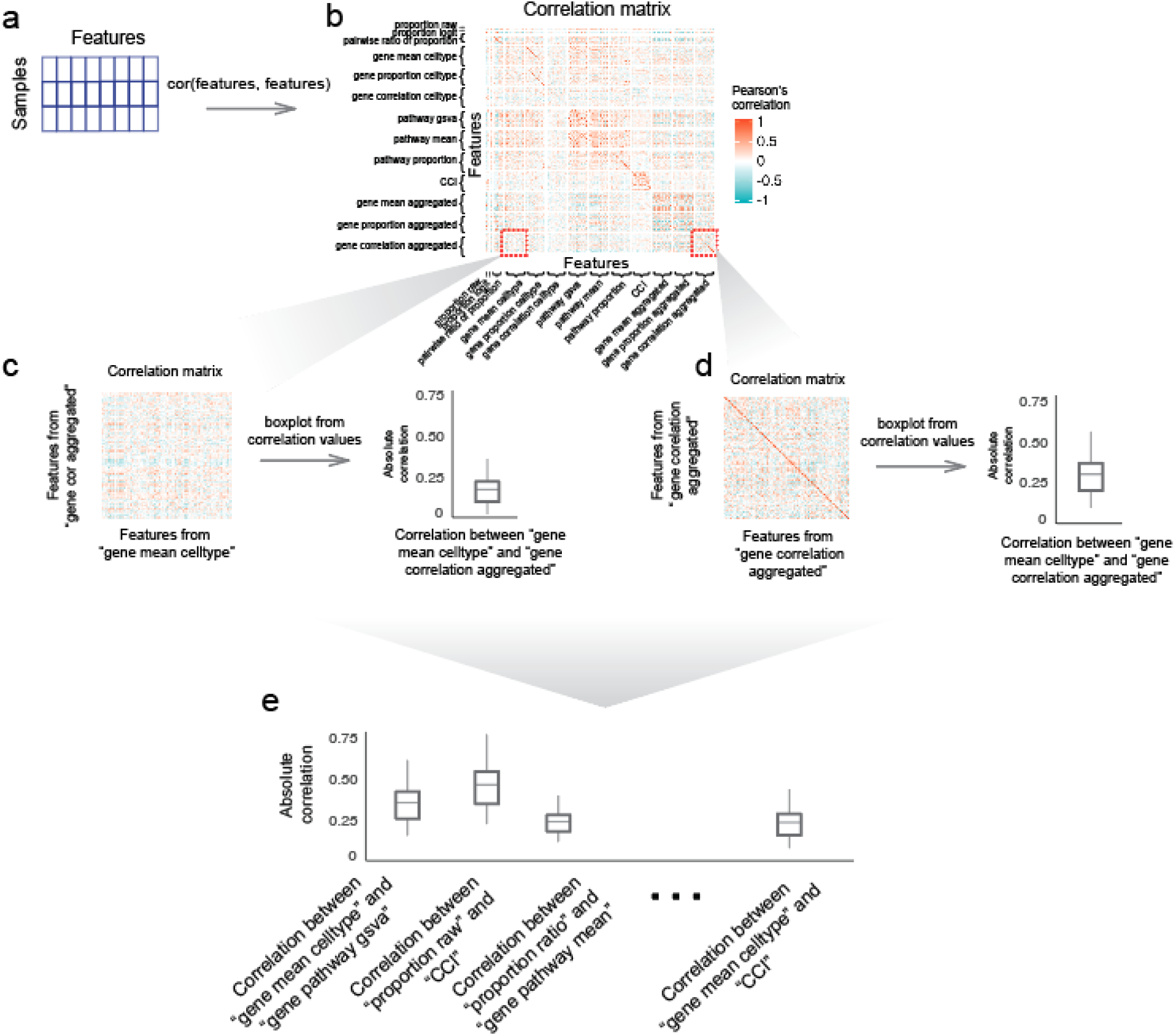
Schematic representation of the calculation of correlation between feature types. **a** First, for a given dataset, the features from all feature types are created, yielding a samples by features matrix. Pearson’s correlation is calculated on the features matrix to result in a typical correlation matrix comprising the correlation between the individual features, as shown in **b**. Since each feature is associated with a feature type, we can zoom into a section of the correlation matrix that contains the correlations of features from two feature types. For exampl**e, c** shows the section of correlation matrix, which contains the features from the feature type “gene mean celltype” and “gene correlation aggregated”. A boxplot can then be constructed to summarise the correlations between these two feature types. **d** shows another section of the correlation matrix, which contains the correlations between all the features from the feature type “gene correlation aggregated”. **e** Repeating this across each section of the correlation matrix produces boxplots summarising the correlation between all pairwise combinations of feature types.

**Supplementary Figure 4.**
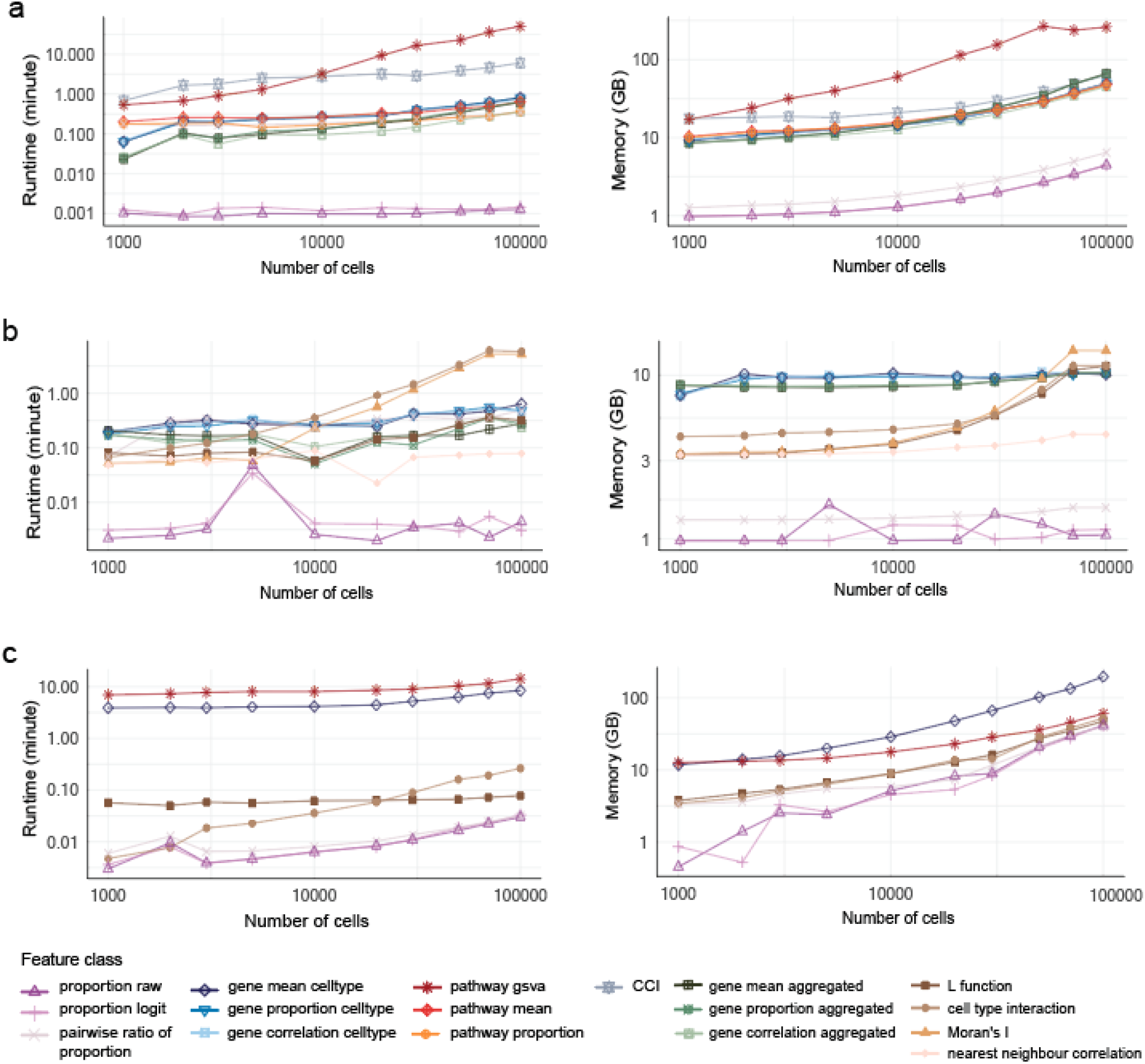
Scalability analysis of feature types. The x and y axes are displayed on a log10 scale in all panels. **a** The runtime and memory usage of feature types benchmarked on subsampled scRNA-seq data. **b** The runtime and memory usage of feature types benchmarked on subsampled spatial proteomics data. **c** The runtime and memory usage of the feature types adapted for the spot-based data, evaluated using spatial transcriptomics data.

**Supplementary Figure 5.**
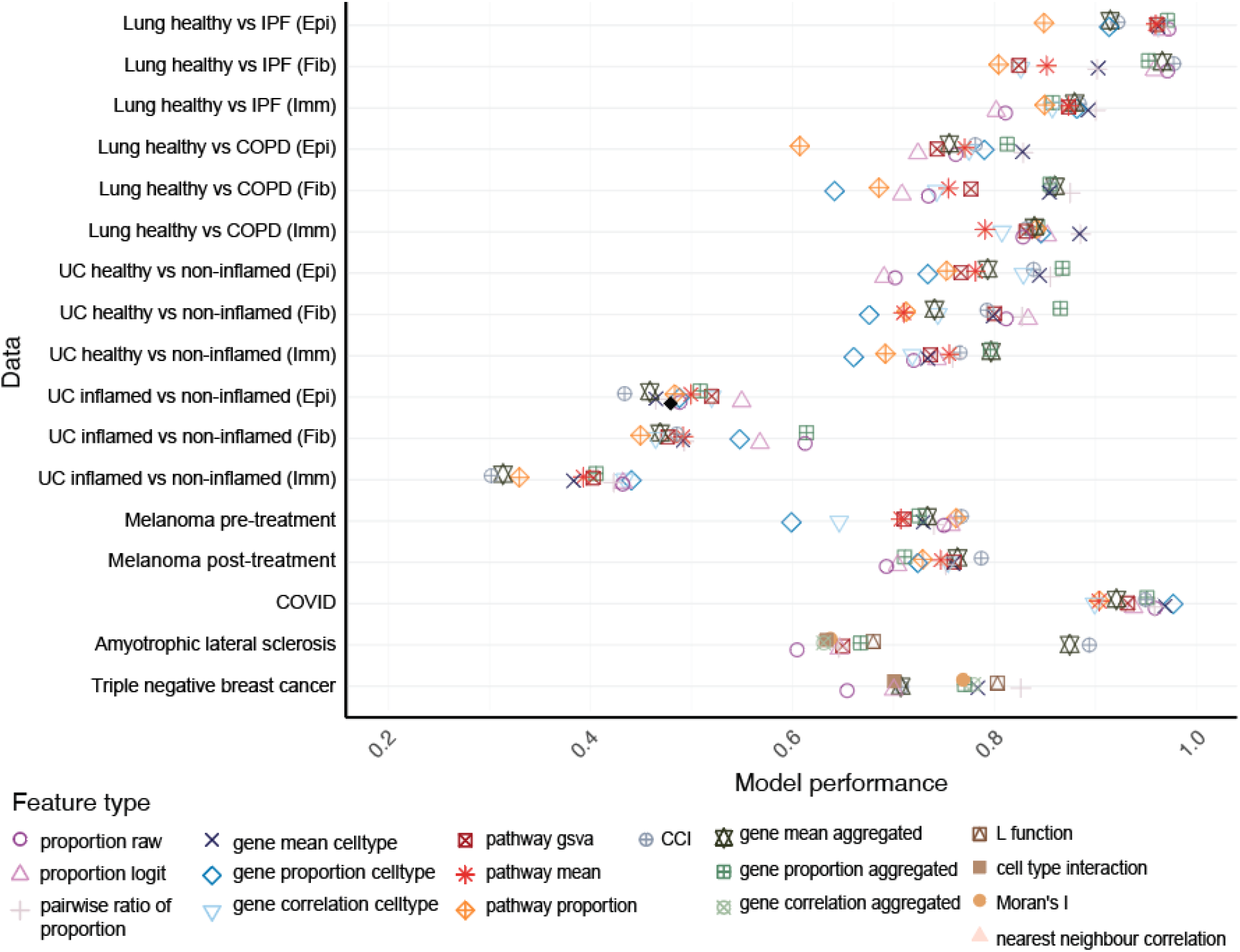
Model performance of each feature class on all datasets. For datasets with disease outcome, random forest was used and model performance was evaluated in terms of F1 score. For the dataset “Triple negative breast cancer” with survival outcome, cox proportional-hazard model was used and model performance was evaluated in terms of C-index. Each point represents the average from 50 cross-validation models.

**Supplementary Figure 6.**
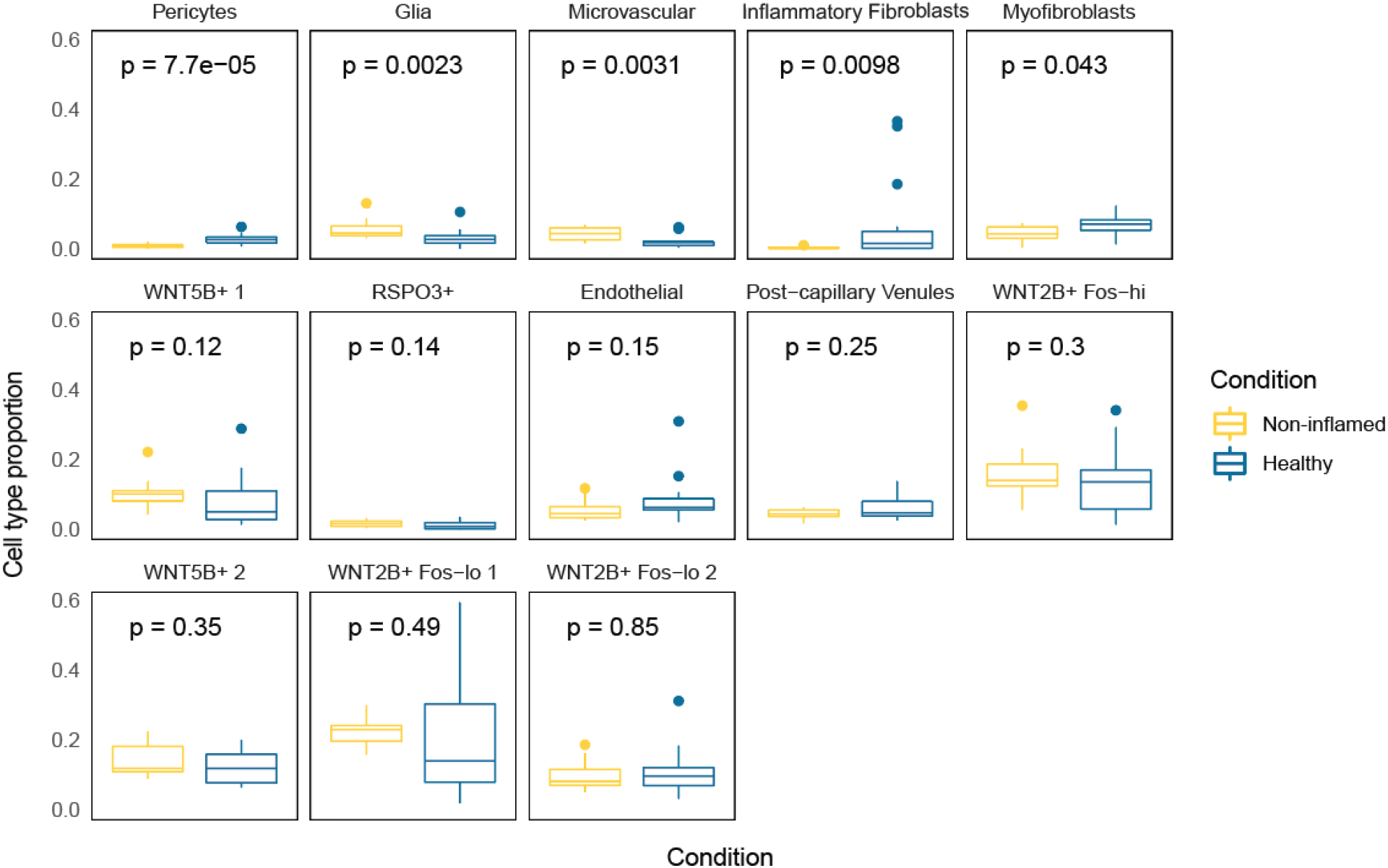
Cell type proportion of the patients in the “UC healthy vs non - inflamed (Fib)” dataset. Wilcoxon test was performed on each cell type to compare the cell type proportion between the non-inflamed and healthy samples.

### Supplementary Tables

**Supplementary Table 1.**
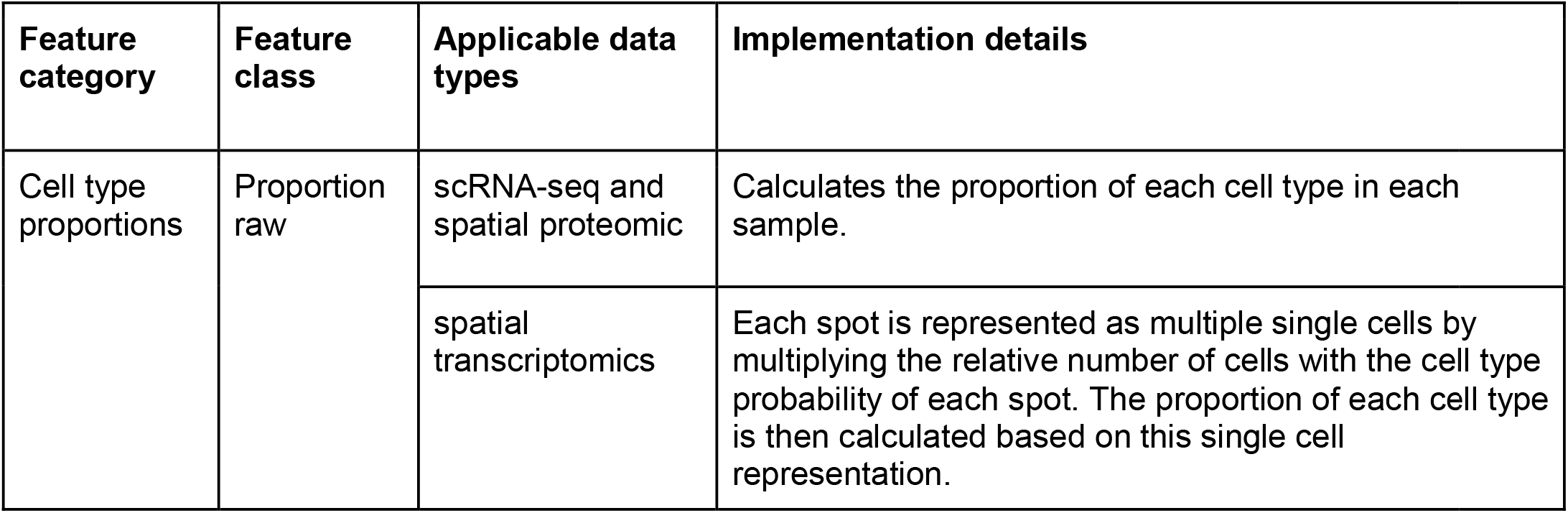
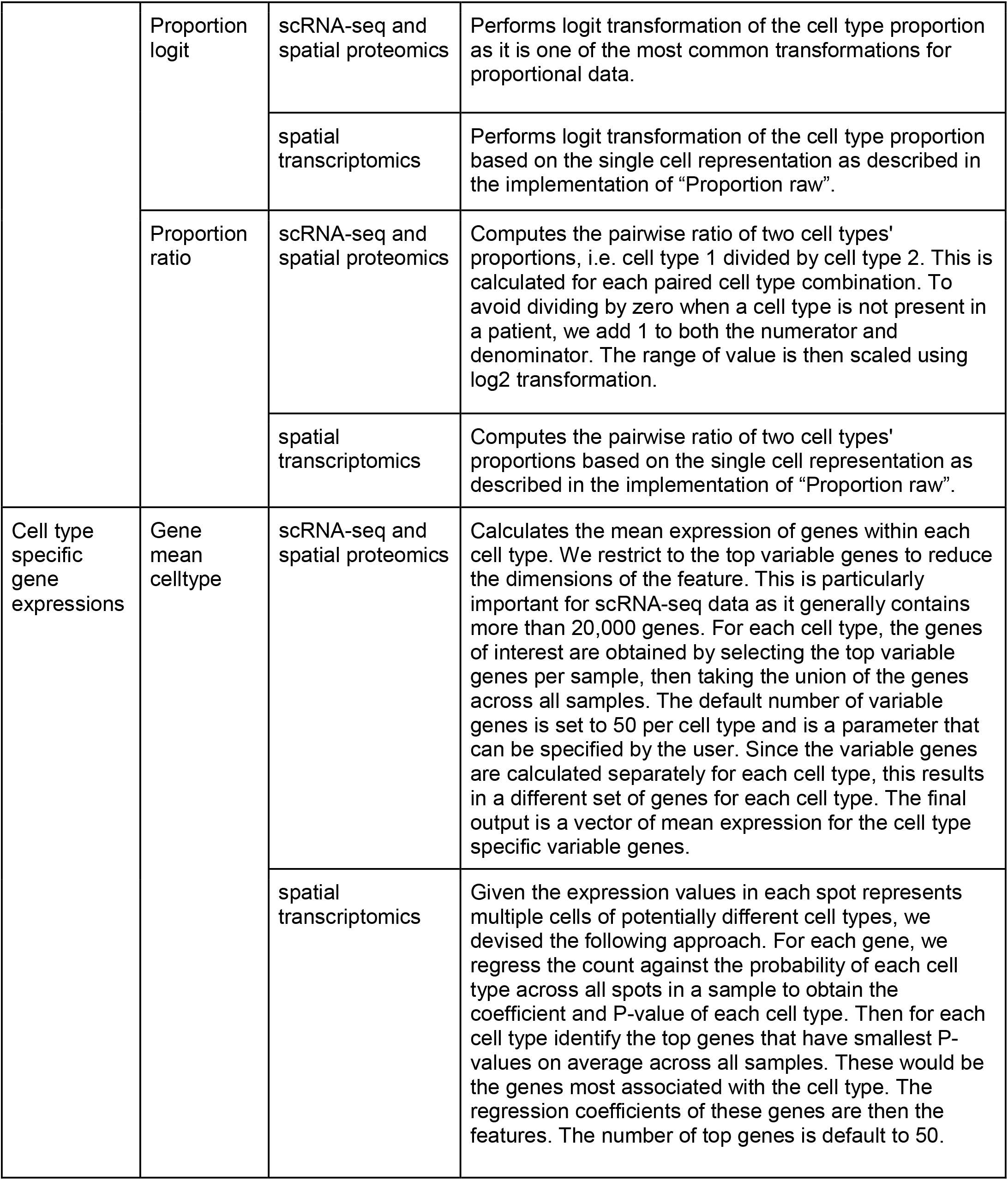
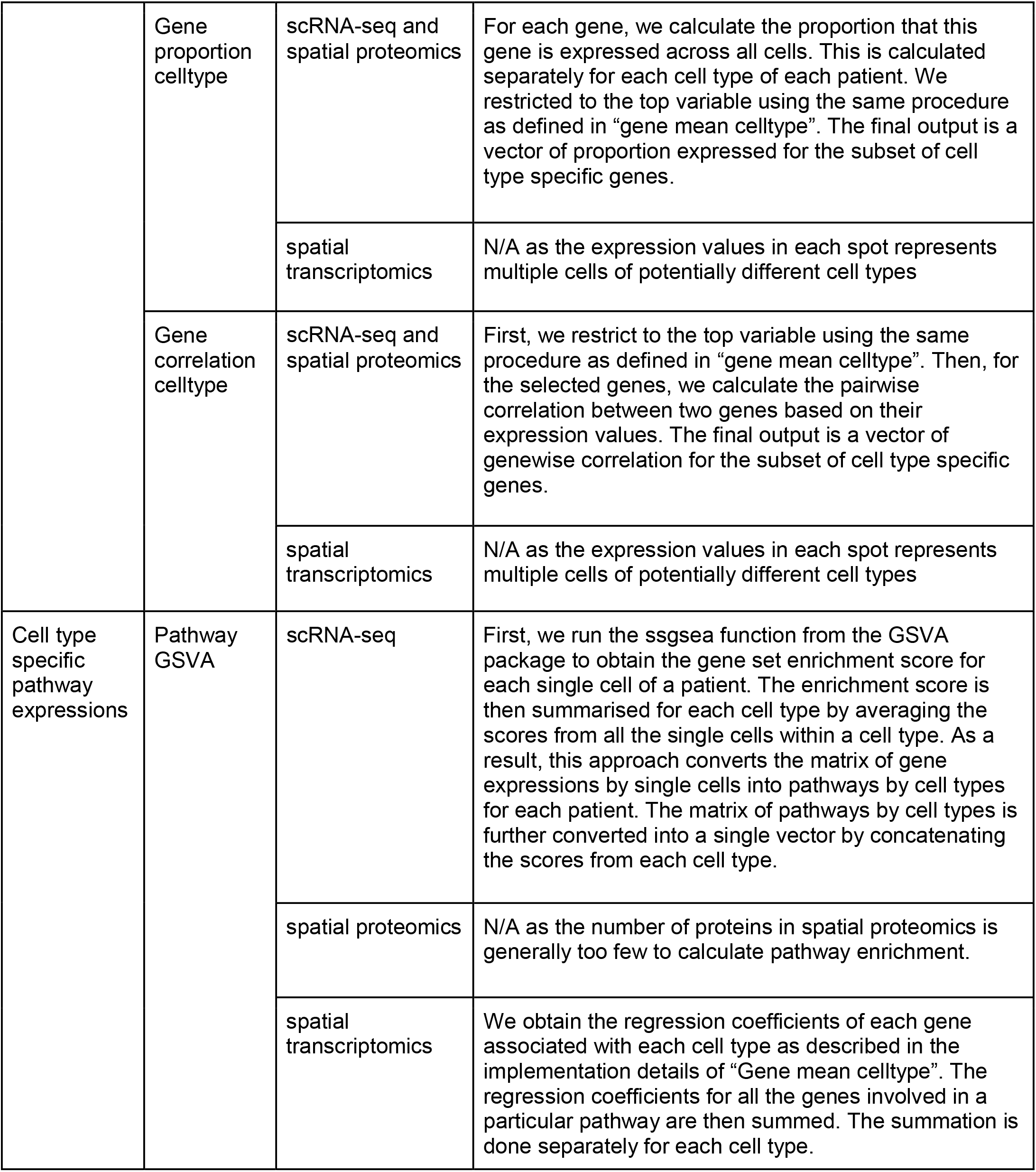
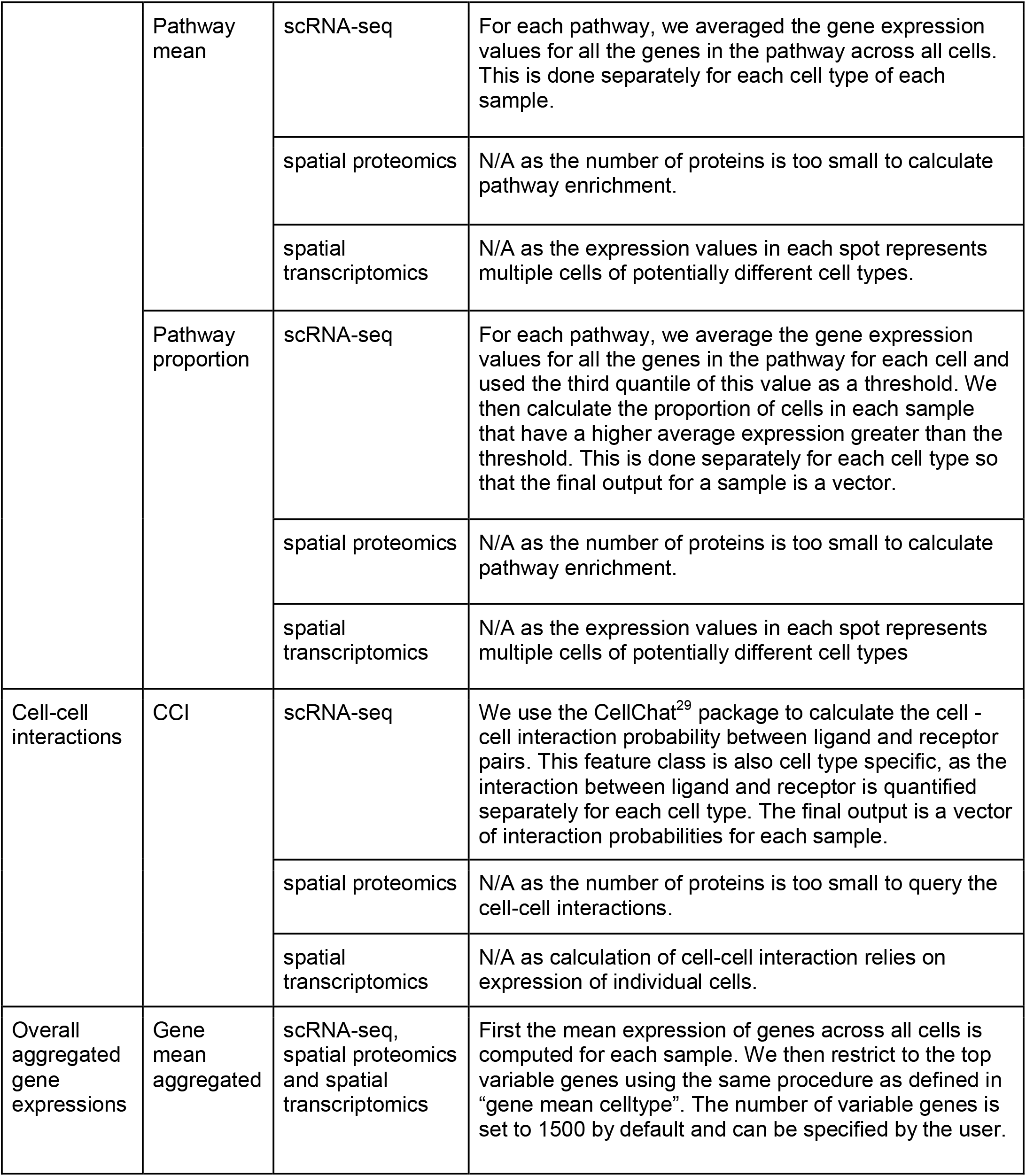
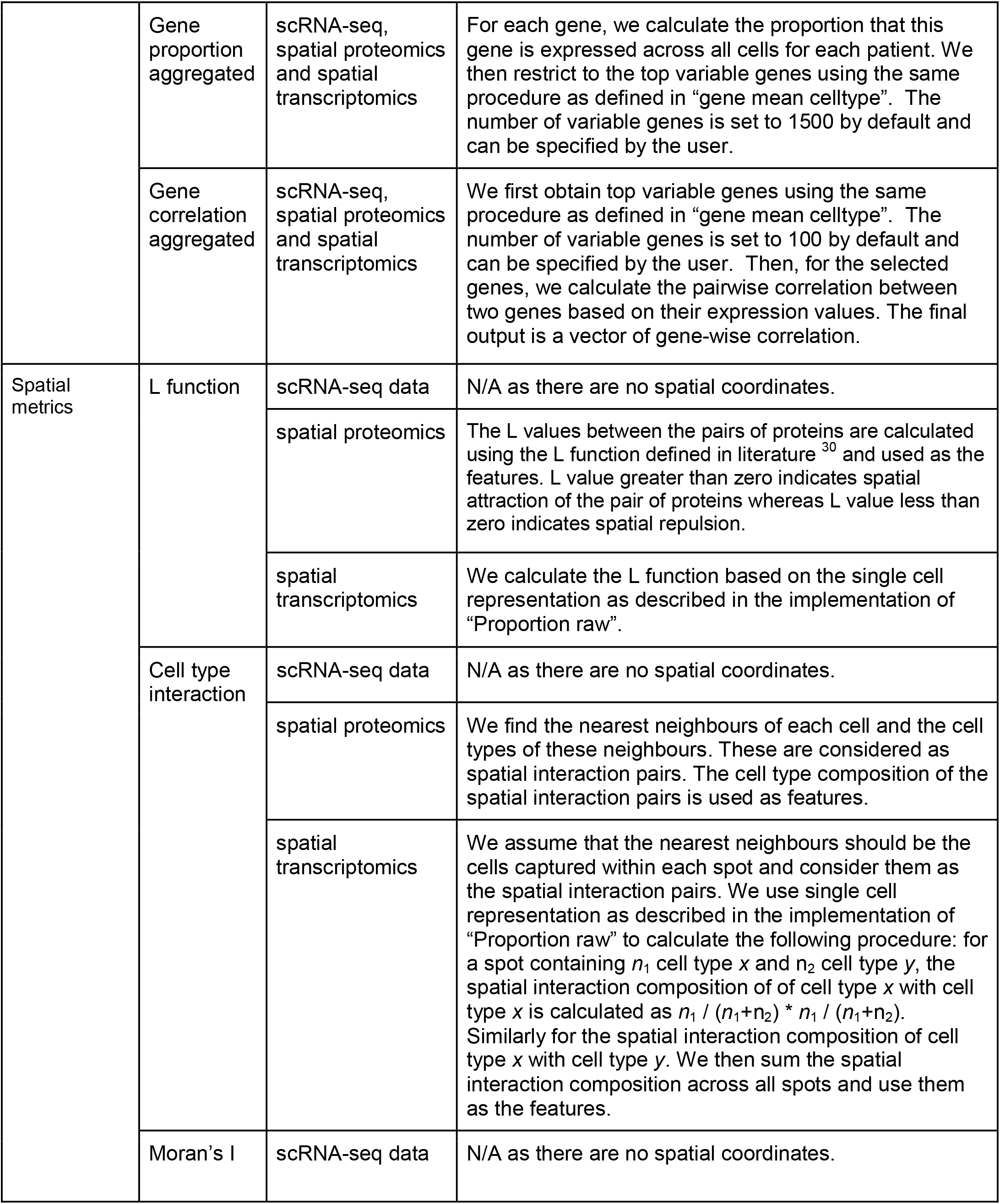
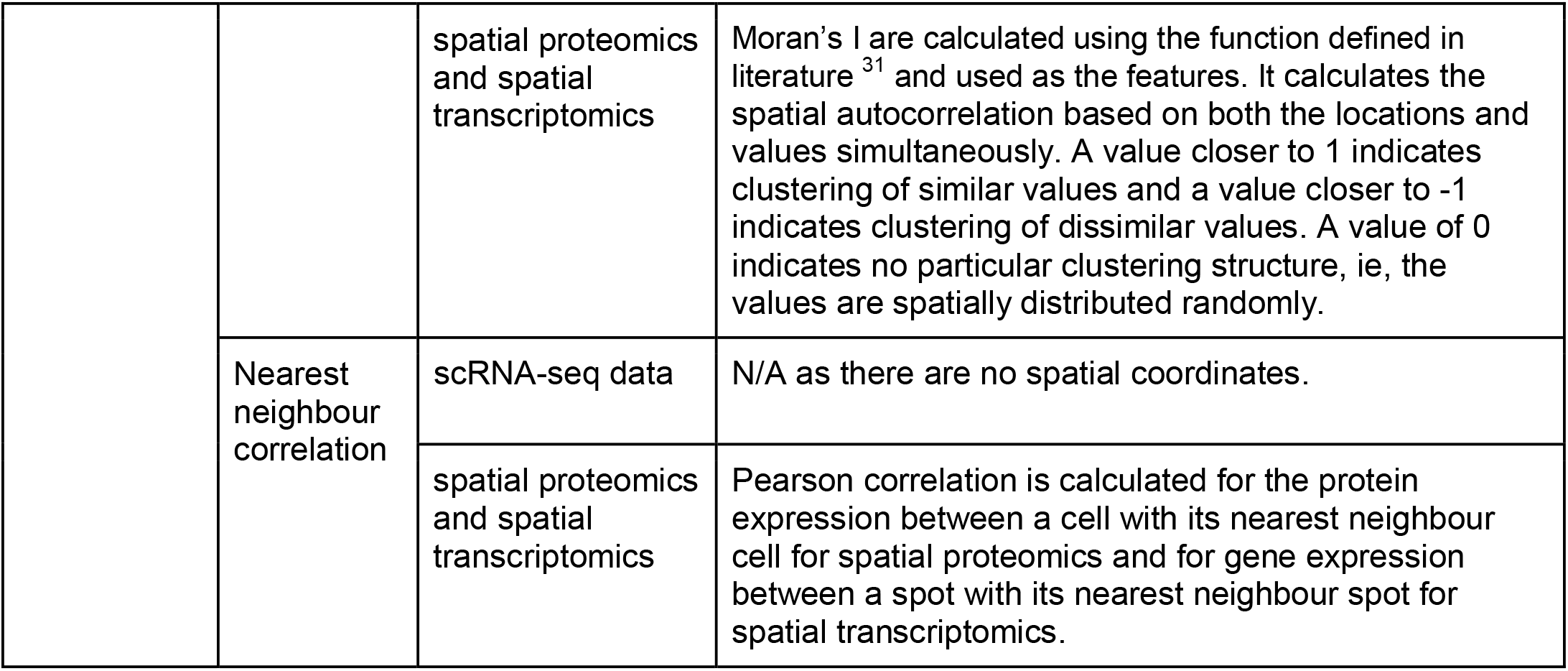

## Appendix

## Acknowledgement

The authors would like to thank all their colleagues, particularly at The University of Sydney, School of Mathematics and Statistics, for their intellectual engagement and constructive feedback. This study was made possible in part by the Australian Research Council Discovery Project Grant (DP170100654) to J.Y.H.Y. and P.Y.; Discovery Early Career Researcher Award (DE170100759) and Australia National Health and Medical Research Council (NHMRC) Investigator Grant (APP1173469) to P.Y.; AIR@innoHK programme of the Innovation and Technology Commission of Hong Kong, Australia NHMRC Career Developmental Fellowship (APP1111338) to J.Y.H.Y.; Research Training Program Tuition Fee Offset and University of Sydney Postgraduate Award Stipend Scholarship to Y.C.

## Data availability

All data used in this study are publicly available. The accession links are reported in the Methods section.

## Code availability

scFeatures is publicly available as an R package at https://github.com/SydneyBioX/scFeatures.

